# Endogenous giant viruses contribute to intraspecies genomic variability in the model green alga *Chlamydomonas reinhardtii*

**DOI:** 10.1101/2021.11.30.470594

**Authors:** Mohammad Moniruzzaman, Frank O. Aylward

## Abstract

*Chlamydomonas reinhardtii* is an important eukaryotic alga that has been studied as a model organism for decades. Despite extensive history as a model system, phylogenetic and genetic characteristics of viruses infecting this alga have remained elusive. We analyzed high-throughput genome sequence data of *C. reinhardtii* field isolates, and in six we discovered sequences belonging to endogenous giant viruses that reach up to several hundred kilobases in length. In addition, we have also discovered the entire genome of a closely related giant virus that is endogenized within the genome of *Chlamydomonas incerta*, the closest sequenced phylogenetic relatives of *C. reinhardtii*. Endogenous giant viruses add hundreds of new gene families to the host strains, highlighting their contribution to the pangenome dynamics and inter-strain genomic variability of *C. reinhardtii*. Our findings suggest that the endogenization of giant viruses can have important implications for structuring the population dynamics and ecology of protists in the environment.

## Introduction

*Chlamydomonas reinhardtii* is a widely studied unicellular green alga with a long history as a model organism that dates back to the 1950s (Sasso *et al*., 2018; Salomé & Merchant, 2019). Despite this long history of research, no viruses that infect *C. reinhardtii* have yet been reported, and as a result the diversity of viruses that infect this alga in nature remain unknown. In a recent study, we identified widespread endogenization of “giant viruses” in numerous green algae, which provides evidence of virus-host interactions that take place in nature (Moniruzzaman *et al*., 2020b). These Giant Endogenous Viral Elements (GEVEs) derive from giant viruses within the phylum *Nucleocytoviricota*, which possess large and complex genomes that can reach up to 2.5 Mbp in length (Philippe *et al*., 2013). Giant viruses often encode complex functional repertoires in their genomes that include tRNA synthetases, rhodopsins, cytoskeletal components, histones, and proteins involved in glycolysis, the TCA cycle, and other aspects of central carbon metabolism (Aylward *et al*.). Recent studies have shown that giant viruses are widespread in the environment and infect a wide range of eukaryotic hosts, including green algae (Schulz *et al*., 2020; Moniruzzaman *et al*., 2020a; Endo *et al*., 2020; Meng *et al*., 2021). The complex genomes of giant viruses coupled with their collectively broad host range and ability to endogenize into the genomes of their hosts provides compelling evidence that they may be important vectors of gene transfer in eukaryotes.

In our initial genomic survey of GEVEs we did not find evidence of endogenous giant viruses in the type strain *C. reinhardtii* (CC-503 cw92). Several studies have recently reported draft genomes of *C. reinhardtii* field isolates, however, and in this study we surveyed these strains for evidence of GEVEs. We report that near-complete genomes of giant viruses are present in several field isolates, and our results suggest that *C. reinhardtii* is a host to at least two distinct lineages of giant viruses. These are the first insights into the diversity and genomic complexity of viruses infecting *C. reinhardtii* in nature. We anticipate that this widely-studied green alga will be a valuable model for future studies of virus-host interactions and the mechanistic aspects of giant virus endogenization.

## Results

We analyzed publicly available high-throughput genome sequencing data for 33 wild strains of *C. reinhardtii*. This data was originally generated for population genomic studies of diverse *C. reinhardtii* strains (Flowers *et al*., 2015; Craig *et al*., 2019; Hasan *et al*., 2019). After *de novo* assembly and annotation (see Methods for details), we identified GEVEs in six of the wild strains (Figure 1A,B). In five of these (CC-2936, 2937, 2938, 3268, and GB-66), the GEVEs range from 315-356 Kb in size and harbored all but one *Nucleocytoviricota* hallmark genes, indicating that near-complete genomes of endogenous giant viruses have been retained in these strains (Figure 1B, Dataset S1). In contrast, CC-3061 harbors a GEVE ~113 Kb in size with 5 out of the 10 hallmark genes, indicating that part of the GEVE was lost over the course of evolution (Supplementary Methods, Dataset S1). Moreover, to ensure that GEVEs were not omitted due to assembly issues we also mapped reads from all genome sequencing projects against the GEVEs, and we identified another highly fragmented GEVE in CC-3059 (see Methods). Lastly, we also analyzed the assembled genome of *Chlamydomonas incerta*, a species phylogenetically closest to *C. reinhardtii*, for which a long-read assembled genome has been recently reported (Craig *et al*., 2021). This analysis revealed a GEVE ~475 Kb long which is integrated within a single 592 Kb contig of this alga (Figure 1B).

**Figure 1:**
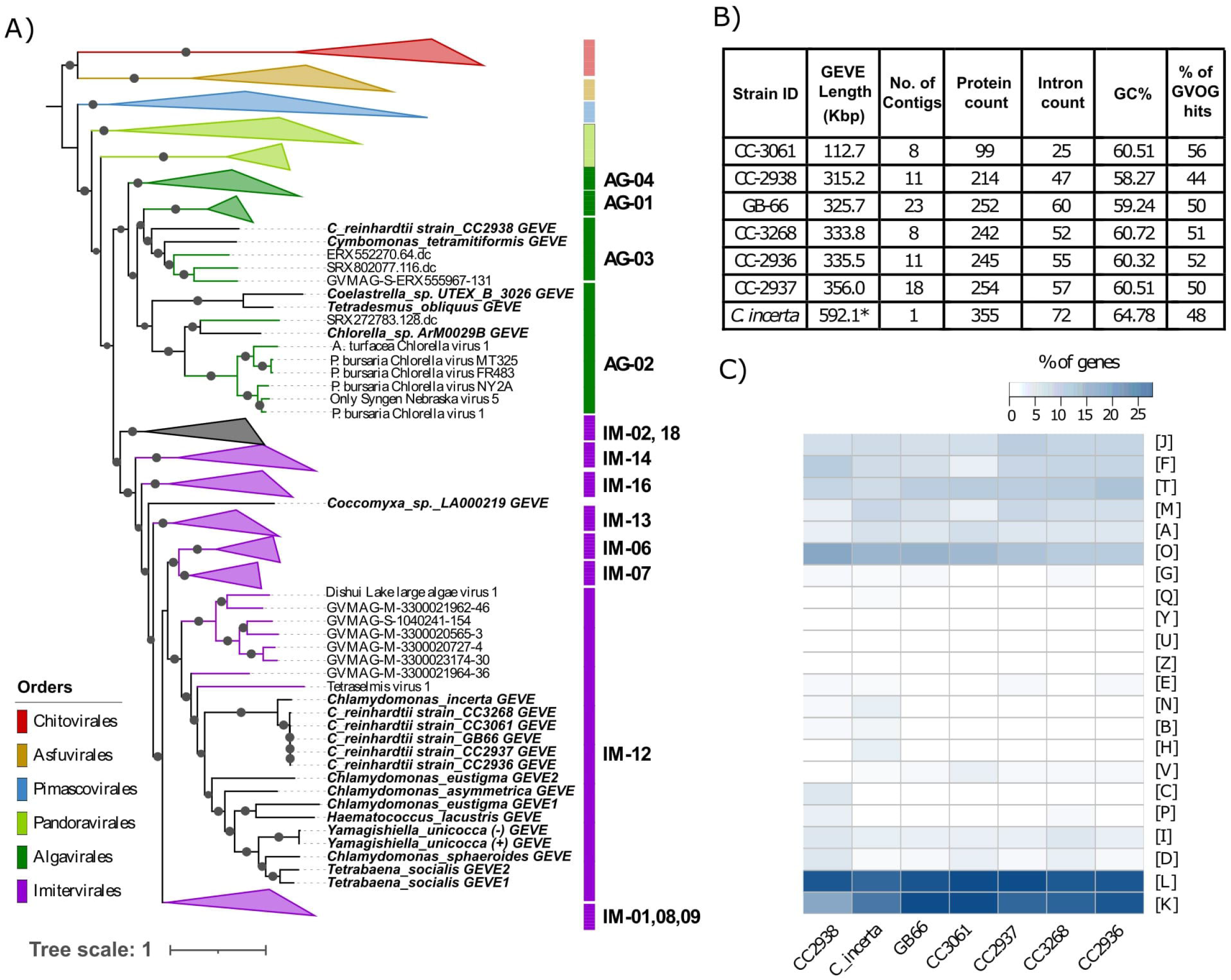
General features and phylogeny of the GEVEs. **A)** Maximum likelihood phylogenetic tree of the GEVEs and representative members from diverse *Nucleocytoviricota* families constructed from a concatenated alignment of seven *Nucleocytoviricota* hallmark genes (see Methods). Individual families within each order are indicated with abbreviations (IM - Imitevirale, AG - Algavirales) followed by family numbers, as specified in Aylward et al, 2019 (Aylward *et al*.). IDs of the GEVEs are indicated in bold-italic. **B)** Basic statistics of the GEVEs present in various field strains of *C. reinhardtii* and the GEVE present in the C. *incerta* genome. **C)** Functional potential of GEVEs as EggNOG categories. Categories of genes are normalized across all the NOG categories except S (function unknown) and R (general function prediction). Raw functional annotations are in Dataset S1. NOG categories: [J] Translation, [F] Nucleotide metabolism, [T] Signal Transduction, [M] Cell wall/membrane biogenesis, [A] RNA processing and modification, [O] Post-translational modification, protein turnover and chamerone, [G] Carbohydrate metabolism, [Q] Secondary structure, [Y] Nuclear structure, [U] Intracellular trafficking and secretion, [Z] Cytoskeleton, [E] Amino acid metabolism, [N] Cell motility, [B] Chromatin structure and dynamics, [H] Coenzyme metabolism, [V] Defense mechanism, [C] Energy production and conversion, [P] Inorganic ion transport and metabolism, [I] Lipid metabolism, [D] Cell cycle control, [L] Replication and repair, [K] Transcription. * *C. incerta* GEVE length includes flanking eukaryotic regions.

Using a newly established taxonomy of *Nucleocytoviricota* (Aylward et al. 2021), we determined the phylogenetic position of the *C. reinhardtii* and *C. incerta* GEVEs and their relationships with other chlorophyte GEVEs that were recently reported (Moniruzzaman *et al*., 2020b) (Figure 1A). Five of the strains harbored GEVEs that formed a cluster within the *Imitervirales* order, consistent with their high pairwise average amino acid identity. The GEVE in *C. incerta* was the closest phylogenetic relative of the *Imitevirales* GEVEs in *C. reinhardtii*, indicating that closely-related giant viruses infect closely related *Chlamydomonas* species in nature. These GEVEs formed a sister clade with the GEVEs present in six other volvocine algae and belonged to the *Imitevirales* family 12 (Figure 1A). Although GEVE contigs could not be recovered from CC-3059, read mapping revealed that this strain also harbors a fragmented *Imitervirales* GEVE (see Methods). In contrast to the GEVEs that could be classified as *Imitervirales*, the GEVE in CC-2938 strain belonged to the *Algavirales* (Figure 1A), indicating that *C. reinhardtii* is infected by multiple phylogenetically distinct lineages of giant viruses in nature.

The coverage of the GEVE contigs was generally similar to those of the host *Chlamydomonas* contigs (see Supplementary Information), consistent with their presence as endogenous elements. The exception was the GEVE in CC-2938, in which two large contigs exhibited the same coverage as those of the host (~8 reads per kilobase per million), while the remaining GEVE contigs had coverage roughly twice that. This unusual pattern may be the product of recent large-scale duplication which recently took place in part of this GEVE. Indeed, recent work on other GEVEs in green algae found that large-scale duplications are common in GEVEs (Moniruzzaman *et al*., 2020b). This would explain why two large contigs with a summed length of 109 kbp retain similar coverage compared to the host contigs, while the rest of the GEVE contigs have roughly double that coverage.

The % GC-content of the *C. reinhardtii* GEVEs ranged from 58.27% (CC-2938) to 60.72% (CC-3268), which is similar to the overall genomic GC content of *C. reinhardtii* (64%) (Merchant *et al*., 2007). Similarly, the GC content of the *C. incerta* GEVE was 64.8%, resembling the overall GC content of the *C. incerta* genome (66%) (Craig *et al*., 2021) (Figure 1B). The GEVEs also contained several predicted spliceosomal introns, ranging from 25 (CC-3061) to 72 (*C. incerta*). Spliceosomal introns are rare in free *Nucleocytoviricota* but have been previously found in GEVEs present in other members of the *Chlorophyta* (Moniruzzaman et al., 2020b). It remains unclear if the relatively high %GC content and spliceosomal introns are features of the viruses themselves, or if the evolution of these features evolved after endogenization. In addition, the GEVE in *C. incerta* was flanked by highly repetitive regions on both ends (Figure 2A). The repetitive region at the 5’-end harbors several reverse transcriptases and transposases (Dataset S1). These regions also have higher intron density compared to the GEVE region itself, and lower number of Giant Virus Orthologous Group (GVOG) hits consistent with their eukaryotic provenance (Figure 2A). This suggests that near-complete genomes of giant viruses can integrate within highly repetitive regions of eukaryotic genomes, potentially with the facilitation of transposable elements.

**Figure 2:**
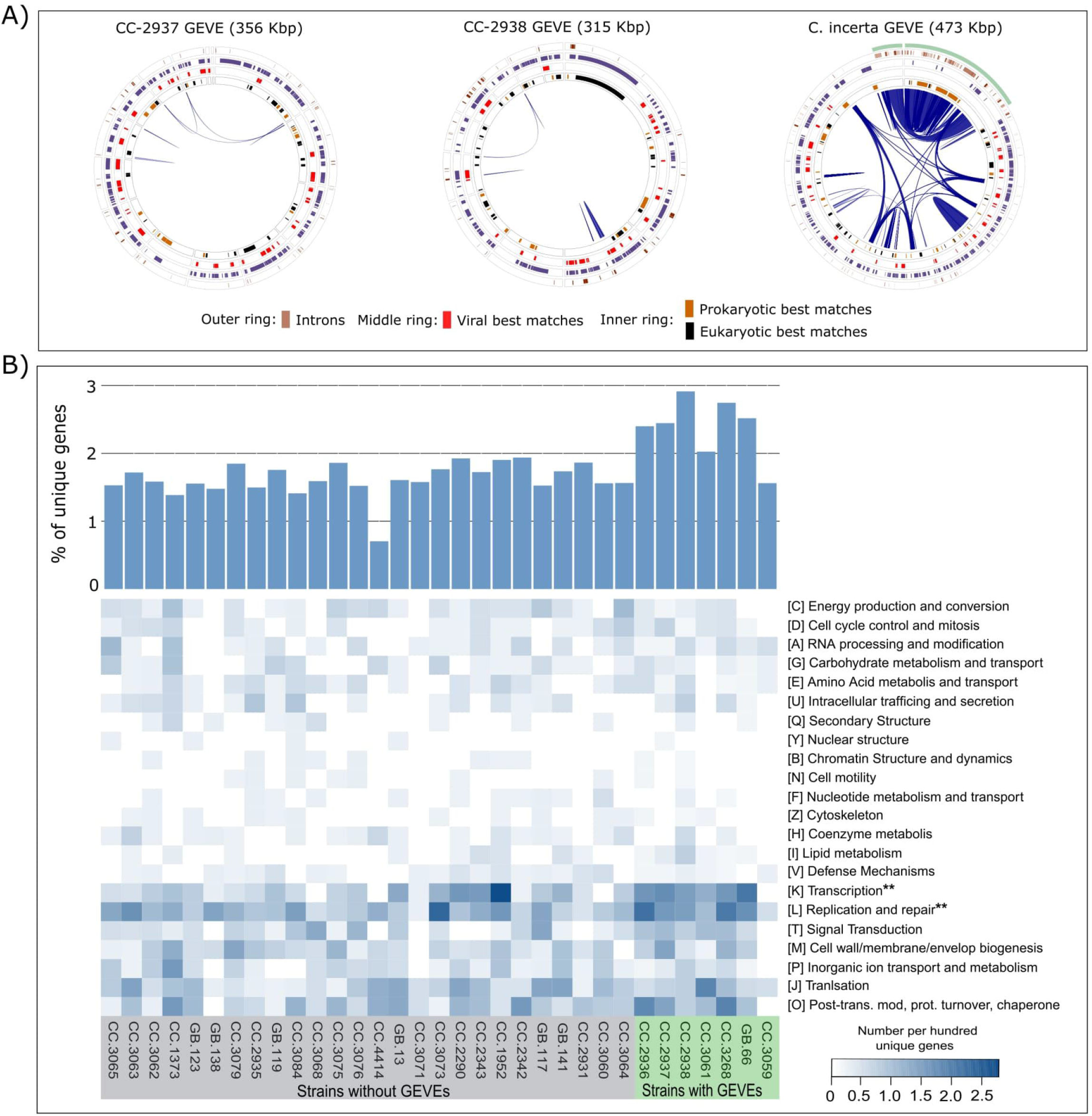
GEVE genomic and functional characteristics. **A)** Circular plots of two representative GEVEs in *C. reinhardtii* and the GEVE present in *C. incerta*. For *C. reinhardtii* one representative *Imitevirales* GEVE (CC-2937) and the Algavirales GEVE (CC-2938) are shown. Circle plots show Giant Virus Orthologous Group (GVOG) hidden Markov model (HMM) hits, spliceosomal introns and the best LAST hit matches (see Supplementary Methods). Internal blue links delineate the duplicated regions. The eukaryotic regions flanking the *C. incerta* GEVE are delineated with light blue stripes. **B)** Unique genes in the field strains of *C. reinhardtii* compared to the reference strain CC-503. The heatmap represents % of unique genes that can be classified in different EggNOG categories (except category [R] - general function prediction and [S] - function unknown). Categories marked with ‘**’ are significantly overrepresented in the GEVE-containing strains compared to those without GEVEs. The bar plot on top of the heatmap represents % of unique genes in each strain. GEVE-containing strains have significantly higher percentages of unique genes compared to the strains without GEVEs.

The GEVEs in *C. reinhardtii* encoded 99 (CC-3061) to 254 (CC-2937) genes, while the *C. incerta* GEVE encoded 355 genes. Most of the genes were shared among the *Imitervirales C. reinhardtii* GEVEs, consistent with their high average amino acid identity (AAI) to each other (>98.5% in all cases, Dataset S1). These GEVEs also shared a high number of orthogroups with the *C. incerta* GEVE (Dataset S1). In contrast, only a few orthogroups were shared between the *Imitevirales and the Algavirales* GEVEs consistent with the large phylogenetic distance between these lineages. Between ~44-55% of the genes in the *C. reinhardtii* and *C. incerta* GEVEs have matches to Giant Virus Orthologous Groups (GVOGs), confirming their viral provenance (Figure 1B). In addition, different genes in these regions have best matches to giant viruses, bacteria, and eukaryotes, which is a common feature of *Nucleocytoviricota* members given the diverse phylogenetic origin of the genes in these viruses (Filée *et al*., 2008) (Figure 2A). Based on the Cluster of Orthologous Group (COG) annotations, a high proportion of the GEVE genes are involved in transcription, and DNA replication and repair; however, genes encoding translation, nucleotide metabolism and transport, signal transduction, and posttranslational modification were also present, consistent with the diverse functional potential encoded by numerous *Nucleocytoviricota* (Figure 1C).

A previous study has shown that several field strains of *C. reinhardtii* harbor many genes that are absent in the reference genome (Flowers *et al*., 2015), which were possibly acquired from diverse sources. To quantify the amount of novel genetic material contributed by giant viruses to *C. reinhardtii*, we estimated the number of unique gene families in the analyzed *C. reinhardtii* field strains that are absent in the reference strain CC-503. On average ~1.78% of the genes in the field strains were unique compared to the reference strain (Figure 2B). Moreover, the GEVE-harboring field strains have significantly enriched in novel genes compared to those without GEVEs (Two-sided Man-Whitney U-test p-value <0.05, Figure 2B). These results suggest that endogenization of giant viruses is an important contributor to inter-strain genomic variability in *C. reinhardtii*. Recent studies have highlighted the importance of horizontal gene transfer (HGT) in structuring the pangenome of diverse eukaryotes (Fan *et al*., 2020; Sibbald *et al*., 2020), and genes originating from endogenous *Nucleocytoviricota* were found to shape the genomes of many algal lineages, including members of the Chlorophyta (Moniruzzaman *et al*., 2020b; Nelson *et al*., 2021). Compared to the GEVE-free strains, GEVE-containing strains harbored a significantly higher proportion of genes from two COG categories including Transcription, and Replication and Repair (Two-sided Mann-Whitney U test p-value <0.05) (Figure 2B). All together, these GEVEs contributed many genes with known functions, including glycosyltransferases, proteins involved in DNA repair, oxidative stress, and heat shock regulation (Dataset S1).

A recent comparative genomic analysis of *C. reinhardtii* analyzed the population structure of this alga by comparing numerous field strains (Craig *et al*., 2019). Interestingly, we found *Imitervirales* GEVEs in both North America populations 1 and 2 (NA1 and NA2, respectively), and in both cases the GEVE-harboring strains are members of populations that include strains for which GEVEs could not be detected. Indeed, strains CC-2931, CC-2932, and CC-3268 were all isolated from the same garden in North Carolina, yet a GEVE could only be detected in CC-3268. This patchwork distribution of the *Imitervirales* GEVEs within *C. reinhardtii* populations suggests that they are the product of independent endogenization events rather than a single event in their shared evolutionary history. Moreover, the *Imitervirales* GEVEs we identified here fall within the same clade as most of the GEVEs we previously identified in other green algae. The prevalence of GEVEs within a particular lineage, together with their patchwork distribution across *C. reinhardtii* strains in the same population, suggests that GEVEs are the product of an active endogenization mechanism that takes place over short timescales rather than “accidental” endogenization that may result from illegitimate recombination that occurs during infection.

## Discussion

While much work remains to elucidate the role of GEVEs in shaping the ecological and evolutionary dynamics of *C. reinhardtii*, several possibilities remain open. Some genes contributed by the GEVEs could be potentially co-opted by the host, leading to changes in certain phenotypes compared to closely related strains without GEVEs. Strain-specific endogenization can also potentially lead to intraspecific variations in chromosome structure, partly mediated by the GEVE-encoded mobile elements (Filée, 2018). Finally, it is also possible that some of these GEVE-loci can produce siRNAs that might participate in antiviral defense, and similar phenomena has been suggested for the virus-like loci in the genome of moss (*Physcomitrella patens*) (Lang *et al*., 2018). Recent studies on the large-scale endogenization of giant viruses into diverse green algal genomes suggest that interactions between giant viruses and their algal hosts frequently shape eukaryotic genome evolution (Moniruzzaman *et al*.) and leads to the introduction of large quantities of novel genetic material. Our results indicate that these endogenization events can lead to genomic variability not only between algal species, but also between strains within the same population. Results reported in this study advance our understanding of how giant viruses shape the genome evolution of their hosts, while also expanding the scope of *C. reinhardtii* as a model organism to study the evolutionary fate and consequences of giant virus endogenization.

## Methods

All methods and relevant citations are available in the ‘Supplementary Information’ file.

## Data and Code availability

Dataset S1 contains information regarding the raw data source, GEVE functional annotations, hallmark gene distribution in each GEVE and coverage information of the partial GEVE in CC-3061.

All the GEVE fasta files, unique gene fasta in each of the strains and their annotations, and concatenated alignment file used to build the phylogenetic tree in Figure 1 are available in Zenodo: https://zenodo.org/record/4958215

Code and instructions for ViralRecall v2.0 and NCLDV marker search scripts are available at: github.com/faylward.

## Acknowledgements

We acknowledge the use of the Virginia Tech Advanced Research Computing Center for bioinformatic analyses performed in this study. This work was supported by grants from the Institute for Critical Technology and Applied Science and the NSF (IIBR-1918271) and a Simons Early Career Award in Marine Microbial Ecology and Evolution to F.O.A.

