## Supplementary Figure 1 for "Endogenous giant viruses contribute to intraspecies genomic variability in the model green alga *Chlamydomonas reinhardtii*"

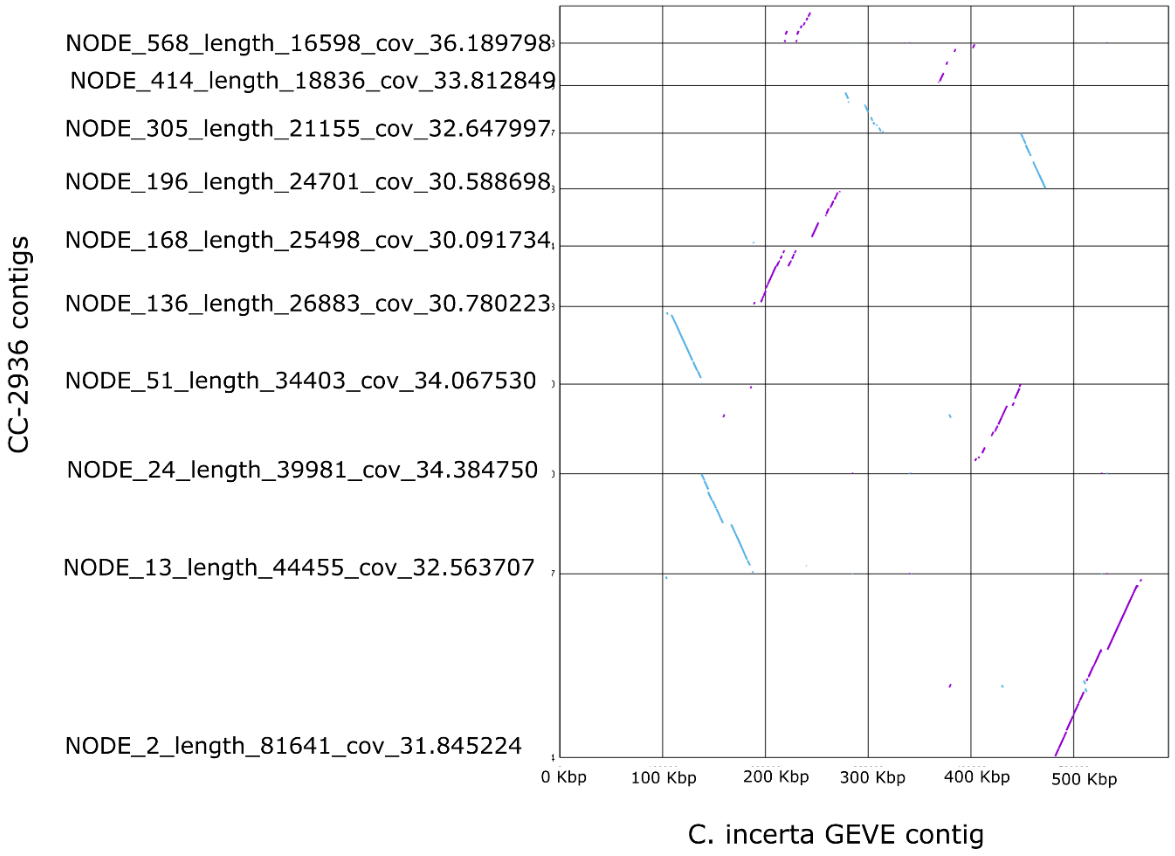


Synteny plots among the final set of ViralRecall screened contigs of the GEVE in CC-2936 against the *C. incerta* GEVE contig (see Methods for details). Amino acid level synteny was calculated using 'promer' implemented in the Nucmer package.

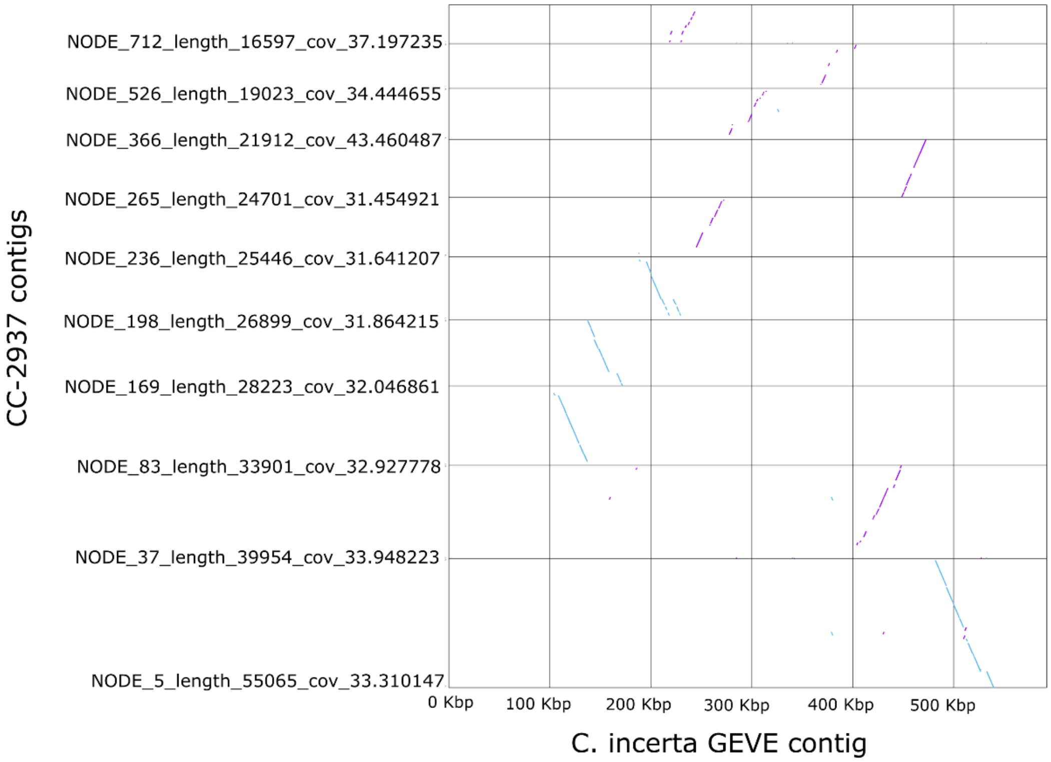


Synteny plots among the final set of ViralRecall screened contigs of the GEVE in CC-2937 against the *C. incerta* GEVE contig (see Methods for details). Amino acid level synteny was calculated using ‘promer’ implemented in the Nucmer package.

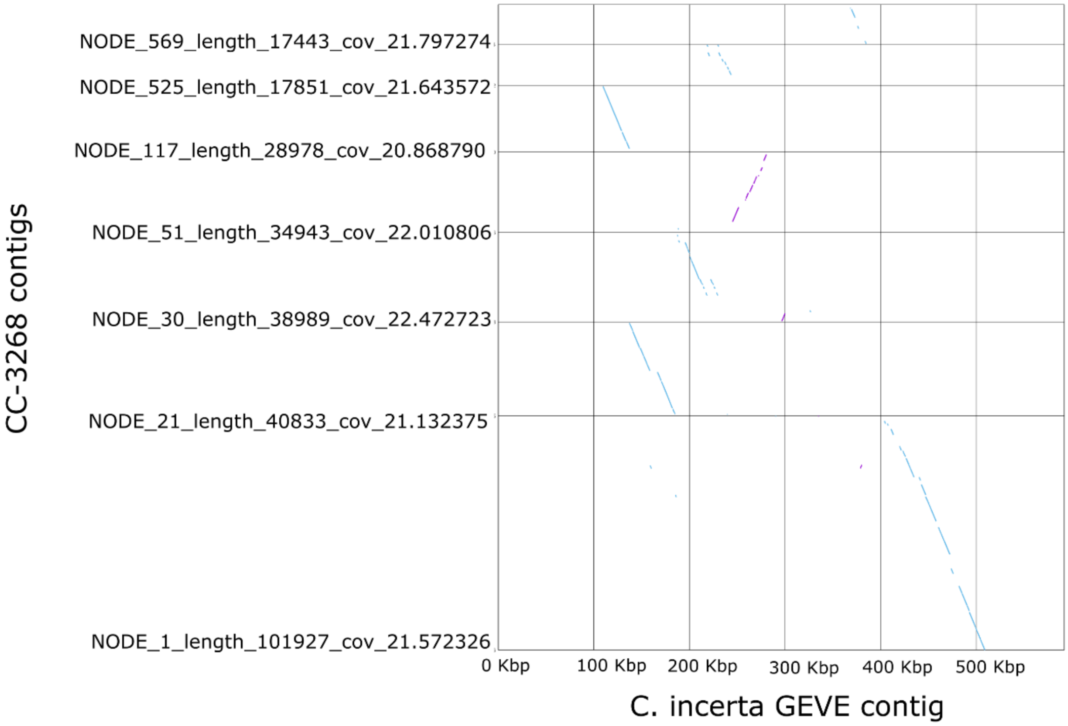


Synteny plots among the final set of ViralRecall screened contigs of the GEVE in CC-3268 against the *C. incerta* GEVE contig (see Methods for details). Amino acid level synteny was calculated using ‘promer’ implemented in the Nucmer package.

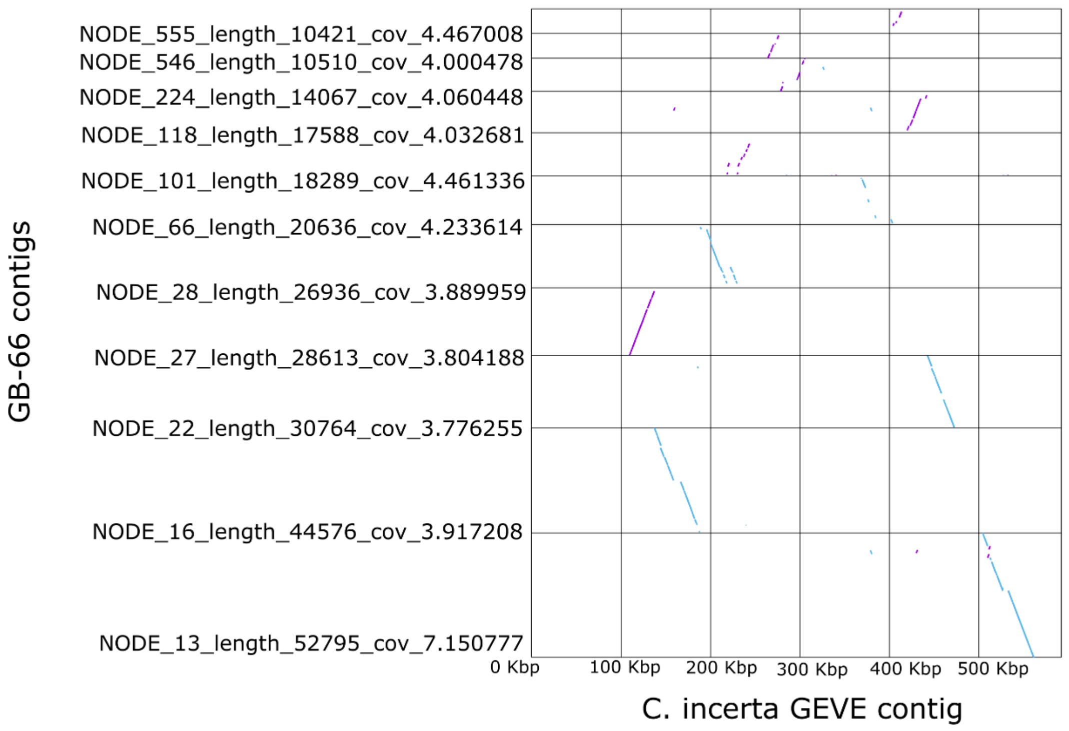


Synteny plots among the final set of ViralRecall screened contigs of the GEVE in GB-66 against the *C. incerta* GEVE contig (see Methods for details). Amino acid level synteny was calculated using 'promer' implemented in the Nucmer package.

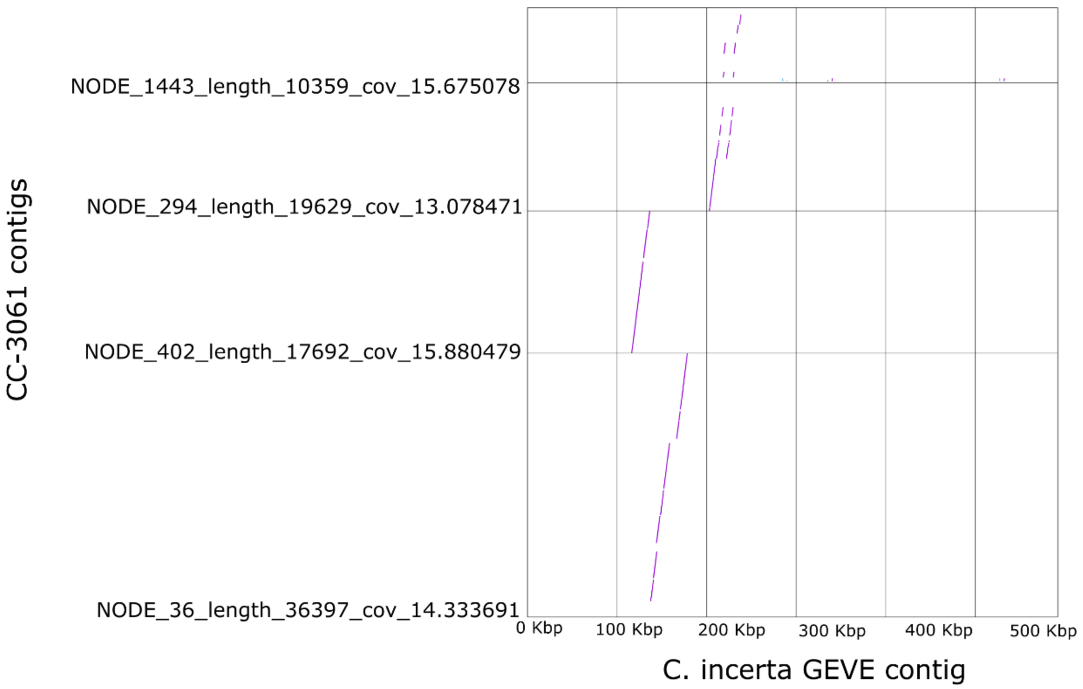


Synteny plots among the final set of ViralRecall screened contigs of the GEVE in CC-3061 against the *C. incerta* GEVE contig (see Methods for details). Amino acid level synteny was calculated using ‘promer’ implemented in the Nucmer package.
