## Supplementary figures and images for "Endogenous giant viruses contribute to intraspecies genomic variability in the model green alga *Chlamydomonas reinhardtii*"

### Supplementary Figure 2

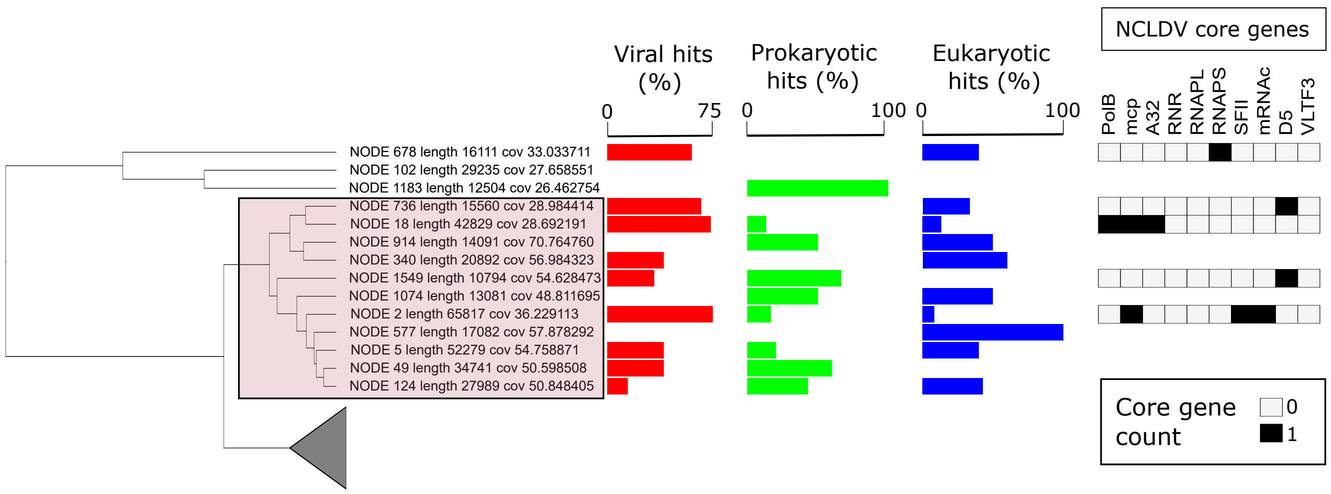

### Supplementary Figure 3

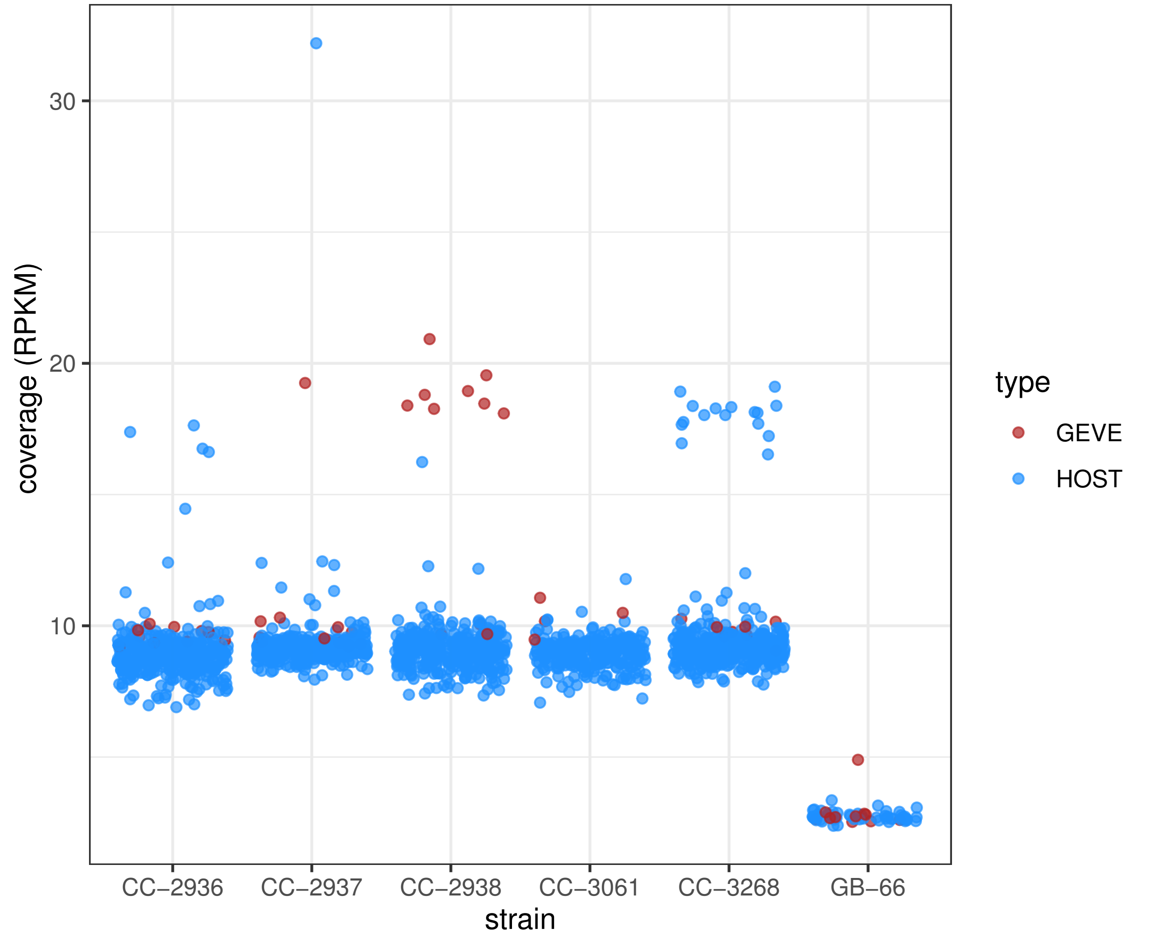
