## Supplementary Information for "Endogenous giant viruses contribute to intraspecies genomic variability in the model green alga *Chlamydomonas reinhardtii*"

Mohammad Moniruzzaman*

Frank O. Aylward*

**This PDF file includes:**

Legends for Supplementary Figure 1, 2, 3, 4, and Datasets S1

SI References

**Supplementary Methods**

**Raw sequence data and genome assembly:** We investigated paired-end illumina sequence data from 33 wild strains that were analyzed in three different studies(1–3). Illumina sequence read libraries were downloaded from NCBI SRA (see Dataset S1). Data from 27 libraries were assembled using SPAdes v3.13.1 (4) (parameters: --meta). For 6 of the libraries (CC-3060, CC-3062, CC-3063, CC-3064, CC-3065, and CC-3073), SPAdes assembler failed as it required more memory than was available on our computing nodes. We assembled these libraries using MEGAHIT (5) with default parameters following quality trimming using TrimGalore (https://github.com/FelixKrueger/TrimGalore) (parameters: --length 36, --stringency 1, -q 5).

**Hybrid gene prediction:** For predicting genes on the final set of GEVE contigs, a hybrid gene prediction approach was taken, based on an approach we developed previously (6). Specifically, we first predicted genes using webAUGUSTUS (7) (http://bioinf.uni-greifswald.de/webaugustus/) and the *Chlamydomonas reinhardtii* training model on the whole assembled genomes of the *C. reinhardtii* strains. For the GEVE harboring strains, we also predicted genes using Prodigal v.2.6.3 (8), which is widely used to predict genes in both prokaryotes and diverse viruses, including NCLDVs. For the GEVE contigs, we retained all the gene and intron predictions by webAUGUSTUS, and also retained the prodigal predicted genes only if they did not overlap with the gene boundaries predicted by webAUGUSTUS. This hybrid approach allowed us to leverage both prediction strategies, as we previously found that some viral genes can be missed by webAUGUSTUS, but were predicted by Prodigal in these regions (6).

**Curation of GEVE contigs:** We identified the preliminary candidate viral contigs from each assembled *C. reinhardtii* strains and *C. incerta* assembled genome using ViralRecall v.2.0 (Description of ViralRecall, parameters). We identified NCLDV hallmark genes in these contigs using a python script that we previously developed (<https://github.com/faylward/ncldv_markersearch>). After identifying the NCLDV hallmark genes in these contig sets, we performed preliminary phylogenies using the DNA polymerase gene which revealed that the endogenous viruses in five of the *C. reinhardtii* strains are highly similar and belongs to the *Imitevirales*, whereas one of these strains harbored endogenous giant virus from the *Algavirales* group. The *C. incerta* GEVE, which was found to be endogenized in its entirety on a large contig, was found to be a close relative of the endogenous *Imitevirales* members from *C. reinhardtii* strains (Figure 1). The 5` and 3’ flanking regions of the *C. incerta* GEVEs (~95 Kb and ~22 Kb, respectively) harbored features characteristic of eukaryotic genomes, specifically, large repetitive regions that have comparatively higher intron density and low number of GVOG hits. The 5’ region also harbored a KDZ transposase (Pfam: 18758), and two copies of zinc-binding regions associated with reverse transcriptases (Pfam: 13966) (Dataset S1). Based on this evidence, we defined the *C. incerta* GEVE to be ~475 Kbp bordered by these two flanking eukaryotic regions.

After determining the phylogenetic provenance of the endogenous viruses in each of these strains, we screened all the contigs detected by ViralRecall v2.0 to remove the contigs that originated from the *C. reinhardtii* reference chromosomal regions. We aligned the contigs to the reference genome chromosomes using Minimap2 (9) and removed contigs that were >90% similar to the reference chromosomes. Contigs that shared >50-90% similarity to the reference genome were manually inspected, and in all cases were found to encode repetitive protein domains of diverse functions. It is possible that these regions originated from the host genome through possible duplication in different strains, and we excluded these contigs from subsequent analyses.

Following these steps, using the remaining set of contigs we delineated the GEVEs in each of these strains harboring *Imitevirales* GEVEs. Given the phylogenetic proximity of the *C. incerta* GEVE and its contiguous assembly in one large contig, we used this GEVE as a guide to validate the *C. reinhardtii* Imitevirales GEVEs. We aligned these contigs against the *C. incerta* GEVE at amino acid level using the promer tool implemented in MUMMER package (10) to assess the similarity of these contigs to *C. incerta* GEVE, and determined all these contigs to be originating from the same viral genome based on their alignment to this GEVE (Supplementary Figure 1). Given the fragmented nature of assembly of individual libraries, it was possible that some of the viral regions were missed by ViralRecall in one strain, but the same region was detected in a different strain if that region was assembled into a larger contig. Since the GEVEs in the *Imitevirales* family are highly similar, we cross-referenced these confirmed viral contigs between libraries to detect smaller contigs that were otherwise missed by ViralRecal in one library but were detected in another. These steps ensured maximum recovery of the viral regions from each library and allowed for a better estimation of the GEVE size and comparative analysis between GEVEs.

To define the *Algavirales* GEVE in CC-2938, we performed a hierarchical clustering of the tetranucleotide frequency of the final ViralRecall screened contigs along with the rest of the contigs from the same host strain (>5kb long). This analysis was performed to ensure that the viral contigs cluster together, and separately from the host contigs, which will be expected based on their distinct viral origin. The results confirmed a cluster of contigs to co-cluster distinctly from the host contigs (Supplementary Figure 2), which was determined to be the *Algavirales* GEVE present in CC-2938.

**Coverage analysis of GEVE and host contigs:** If GEVE contigs were truly endogenous we would expect them to have similar coverage to that of the rest of the *C. reinhardtii* genome. To test this, we compared the coverage of the GEVE and host contigs by mapping reads from each genome onto its assembly. We performed read mapping with CoverM (<https://github.com/wwood/CoverM>) with the parameter “--min-covered-fraction 50”. To ensure that host contigs belonged to *C. reinhardtii* chromosomes, we compared all contigs to the reference 17 chromosomes of *C. reinhardtii* strain CC-502 cw92 mt+ with LAST (default parameters) and retained only contigs with an e-value match of 1e-100. For this analysis we only considered host contigs > 20 kbp in length and GEVE contigs > 10 kbp in length.

**Read mapping to confirm GEVE absence:** In the strains in which we did not identify GEVEs we sought to confirm that their absence was real and not simply due to complications arising from *de novo* assembly. For this we mapped raw sequencing reads from all genomes against the set of GEVE contigs from CC-2938 and CC-2937. These two were chosen because CC-2938 is the sole *Algavirales* GEVE that we found, while CC-2937 was the *Imitervirales* GEVE with the largest assembly recovered. We mapped reads using CoverM (<https://github.com/wwood/CoverM>) with the parameter “--min-covered-fraction 50”. Using this approach we confirmed the absence of GEVEs from all strains except CC-3059, where reads could be mapped to 4 of the 14 reference contigs of the *Imitervirales* GEVE. This suggests that CC-3059 contains a partial GEVE that could not be resolved in the *de novo* assemblies, although in all other cases no GEVE contigs could be recovered.

**Functional annotation:** We predicted function of the protein sequences in each of the GEVEs and the unique genes present in each field strain by searching the proteins against HMM profiles from COG (11), Pfam v. 32 (12), EggNog v. 5.0 (13), eggNOG Viral (13) and VOG (vogdb.org) databases using ‘hmmsearch’ command implemented in HMMER v.3.21(14) with an e-value threshold of <0.00001. Best hit for a protein was evaluated based on the highest bit score to a HMM profile.

**Identification of unique genes in diverse field strains:** For identification of unique genes that are present in different field strains of *C. reinhardtii* but absent in the reference genome (CC-503), we first predicted genes in all these genomes using webAugustus (7) as described in the ‘Hybrid gene prediction’ section. Some strain assemblies contained contigs with coverage >20 times the longest contigs in the assembly, and manual inspection revealed that they likely derived from bacterial contamination. To mitigate the impact of this on our unique gene estimates, for this analysis we did not consider contigs that had coverage greater than one standard deviation above the mean for a given assembly. For the remainder of the contigs, the predicted proteins from the field strains with and without GEVEs were searched against the reference CC-503 proteins (*C. reinhardtii* assembly version 5.5) using ‘Blastp’ (parameters: -max_hsps 1, -max_target_seqs 1). To obtain a conservative estimate of the unique gene families in each field strain, only genes that had no homology at an e-value threshold of 0.001 to the reference proteome were considered. Although we predicted additional genes using Prodigal in the GEVE contigs for GEVE functional analysis and homology searches, for estimating unique genes, we excluded the Prodigal predicted proteins. This was done as we compared results across all field strains - since Prodigal prediction is only relevant for the GEVE contigs, and can not be included for the other contigs in the genome.

**GVOG analysis:** For identifying GEVE genes with similarities to diverse giant viruses, we used a curated Giant Virus Orthologous Group (GVOG) database that we recently constructed from 1,380 quality-checked genomes that include 1,253 metagenome assembled genomes (MAGs) and 127 complete genomes available in culture for *Nucleocytoviricota* members. GVOG are publicly available at (<https://faylward.github.io/GVDB/>) (15). To evaluate hits to the GVOGs, we used ‘hmmsearch’ implemented in HMMER v.3.2.1 with an e-value threshold of <0.00001.

**Duplication and Synteny analysis:** We compared synteny between different GEVE regions using the ‘progressiveMauve’ tool implemented in Mauve package (16). For determining the similarity of the *C. reinhardtii* GEVE contigs to the *C. incerta* GEVE at amino acid level, we used the ‘promer’ tool implemented in MUMMER (10) with the ‘--maxmatch’ option. We estimated the amount of repetitive regions within each GEVE using RECON 1.0.8 (17), with a nucleotide identity of >90%.

**AAI and orthogroup analysis:** AAI between GEVE proteomes was calculated using a custom Python script (https://github.com/faylward/lastp_aai), which carries out pairwise LAST (v. 959) searches (parameters: -m 500) of protein sequences and calculate the average amino acid identities between all possible pairs of genomes (18). Orthogroups of proteins among the GEVEs were calculated using ProteinOrtho v.6.0.6 (19) with default parameters (-identity=25, -cov=50).

**GEVE phylogenies:** For phylogenetic reconstruction of the GEVEs along with known NCLDVs, we used a subset of high-quality genomes recently curated to develop a phylogenomic framework of *Nucleocytoviricota* (15). The GEVEs from this study and a previous study (6) were included. We used a concatenated alignment of a set of 9 core genes as described previously to be ideal for phylogenetic reconstruction of *Nucloecytoviricota* (15). Alignments were generated using Clustal Omega (20) and trimmed with TrimAl (21). The tree was constructed using IQ-TREE v1.6.9 (22) and 1000 ultrafast bootstrap replicates were performed to assess statistical support at the nodes (parameters: -wbt -bb 1000 -m LG+I+G4). The tree was visualized using iTOL (23).

**Homology search:** To identify the best match of the GEVE proteins in diverse domains of life and viruses, we compared the GEVE proteins against a database of NCBI RefSeq v. 99 (24). To this end, we employed LASTAL v. 959 with parameter ‘-m 5000’ for increased sensitivity of homology detection. Before evaluating the best hits, all the hits to Chlorophyta were removed, to avoid self-hits. Taxonomic profile of each best hit was determined by cross-referencing the hits to the NCBI Taxonomy database (25). For this, we used the Python API implemented in the ETE3 Toolkit (26).

**Assessing the partial loss of GEVE in CC-3061:** Total length of the final set of screened GEVE contigs in CC-3061 was ~115 Kb long, which is much smaller than the other *C. reinhardtii* strains harboring GEVEs. This suggests that the GEVE in CC-3061 went through partial loss over the course of genome evolution. However, it is also possible that due to fragmented assemblies obtained from the raw data, some of the GEVE regions failed to assemble and hence were missed by our screening approach. If that’s the case, we should still be able to find the reads corresponding to such small contigs. As the *Imitevirales* GEVEs in the *C. reinhardtii* strains are highly similar to each other, we mapped the raw reads from CC-3061 library to one of the near-complete GEVEs from strain CC-2937. We found that although several of the CC-2937 GEVE contigs had good coverage, other contigs had zero or near-zero coverage, indicating that reads originating from these regions are absent in the CC-3061 library (Coverage values available in Dataset S1). This analysis confirmed that the regions missing in the CC-3061 GEVE are due to partial loss, and not an artefact of lower quality assembly or low sequencing depth.

**Supplementary figure 1:** Synteny plots of the final set of ViralRecall screened contigs of the *C. reinhardtii* GEVEs and the *C. incerta* GEVE contig. Amino acid level synteny was calculated using ‘promer’ implemented in Nucmer package.

**Supplementary figure 2:** Binning of CC-2938 GEVE candidate contigs based on tetranucleotide frequency. Tetranucleotide frequency data was unit variance scaled, followed by average linkage clustering applied on the correlation distance matrix. The cluster of contigs that represent the GEVE is shaded in red. Bar charts show the proportion of best LAST hits to NCLDVs, prokaryotes or eukaryotes on each of the contigs. The collapsed clades represent contigs that potentially originated from host chromosomes and not part of the GEVE.

**Supplementary figure 3.** Coverage of the GEVE and host contigs for the *C. reinhardtii* strains that harbored GEVEs.

**Dataset S1 legend:** Dataset containing functional annotations of the GEVE proteins, Core gene distribution across individual contigs in each of the GEVEs, coverage of the near-complete *Imitevirales* GEVE in CC-2937 from the CC-3061 illumina reads, and references of the raw data used in this study.
